# Msps Governs Acentrosomal Microtubule Assembly and Reactivation of Quiescent Neural Stem Cells

**DOI:** 10.1101/2020.01.24.918227

**Authors:** Qiannan Deng, Ye Sing Tan, Liang Yuh Chew, Hongyan Wang

**Author notes:** Lead contact: Hongyan Wang.

## Abstract

The ability of stem cells to switch between quiescence and proliferation is crucial for tissue homeostasis and regeneration. *Drosophila* quiescent neural stem cells (NSCs) extend a primary cellular protrusion from the cell body prior to their reactivation. However, the structure and function of this protrusion are not well established. In this study, we show that in the primary protrusion of quiescent NSCs microtubules are predominantly acentrosomal and oriented plus-end-out, distal to the cell body. We have identified Mini Spindles (Msps)/XMAP215 as a key regulator of NSC reactivation and acentrosomal microtubule assembly in quiescent NSCs. We show that E-cadherin, a cell adhesion molecule, is localized to NSC-neuropil contact points, in a Msps-dependent manner, and is intrinsically required for NSC reactivation. Our study demonstrates a novel mechanism by which Msps-dependent microtubule assembly in the primary protrusion of quiescent NSCs targets E-cadherin to NSC-neuropil contact sites to promote NSC reactivation. We propose that the neuropil functions as a new niche for promoting NSC reactivation, which may be a general paradigm in mammalian systems.

## INTRODUCTION

The ability of stem cells to switch between quiescence and proliferation is crucial for tissue homeostasis and regeneration. Most neural stem cells (NSCs) that reside in mammalian adult brains are in a mitotically dormant, quiescent state (Doetsch et al., 1999; Morshead et al., 1994). In response to physiological stimuli such as the presence of nutrition and physical exercise, quiescent NSCs (qNSCs) can exit from quiescence and become reactivated to generate new neurons (Fabel and Kempermann, 2008). Conversely, stress, anxiety and old age reduce the proliferation capability of NSCs (Lucassen et al., 2010). Dysregulation of quiescence or reactivation in the nervous system can result in depletion of NSC population or too few differentiated neurons (Cheung and Rando, 2013).

Recently, *Drosophila* NSCs, also known as neuroblasts, have emerged as a powerful model to study the mechanisms underlying NSC quiescence and reactivation *in vivo*. *Drosophila* neural stem cells in the central brain and thoracic ventral nerve cord (VNC) enter into quiescence at the end of embryogenesis and subsequently exit quiescence (also termed reactivation) largely within 24 hours in response to the presence of dietary amino acids (Isshiki et al., 2001; Lai and Doe, 2014b; Tsuji et al., 2008). Their reactivation depends on an evolutionarily conserved insulin/IGF signaling pathway (Britton and Edgar, 1998; Chell and Brand, 2010a; Homem and Knoblich, 2012; Sousa-Nunes et al., 2011a; Truman and Bate, 1988b; Tsuji et al., 2008). Insulin/IGF-like peptides Dilp2 and Dilp6 are secreted by blood-brain-barrier (BBB) glia and activate the insulin/IGF/Akt pathway in the underlying NSCs (Chell and Brand, 2010b; Sousa-Nunes et al., 2011b). Gap junctions in the BBB glia couple metabolic signal with synchronized calcium pulses and insulin secretion, leading to a relatively synchronized reactivation of NSCs (Spéder and Brand, 2014). Mammalian Insulin-like growth factor-1 (IGF-1) and IGF-2 also promote NSC proliferation (Aberg et al., 2003; Mairet-Coello et al., 2009; Yan et al., 2006; Ye et al., 2004). Interestingly, human IGF1R mutations are associated with microcephaly, a neurodevelopmental disorder (Juanes et al., 2015). In the absence of nutrition, the hippo pathway inactivates Yokie to maintain the quiescence of NSCs (Ding et al., 2016; Poon et al., 2016). The hippo pathway is inactivated in response to dietary amino acids and is downregulated by the CRL4-Mahjong E3 ligase complex (Ly et al., 2019). NSC reactivation also requires intrinsic mechanisms involving a transcription factor Prospero, spindle matrix proteins and striatin-interacting phosphatase and kinase (STRIPAK) family proteins (Gil-Ranedo et al., 2019; Lai and Doe, 2014a; Li et al., 2017).

A hallmark of qNSCs in *Drosophila* is the cellular extension(s) attached to the cell body. qNSCs in *Drosophila* extend a primary cellular protrusion towards neuropil and occasionally extend a second but a much shorter protrusion at the opposite side of the cell body (Chell and Brand, 2010b; Truman and Bate, 1988a). The cellular protrusions of qNSCs are removed presumably via retraction prior to their cell cycle re-entry (Chell and Brand, 2010b). Recently, we reported that primary cellular extension of qNSCs is a microtubule - enriched structure (Li et al., 2017). Microtubules are polar filaments with a fast-growing plus-end and a slow growing minus-end. Microtubules have distinct orientations in axons and dendrites of *Drosophila* neurons, with plus ends distal to the cell body (plus-end-out orientation) in axons and opposite orientation in dendrites (Stone et al., 2008). These distinct microtubule orientations are associated with different structures and functional properties of axons and dendrites. However, microtubule orientation in the cellular protrusion of qNSCs is unknown and key microtubule regulators during NSC reactivation have not been identified.

Mini spindles (Msps), a XMAP215/ch-TOG/Msps family protein is a key regulator of microtubule growth in dividing cells (Chen et al., 2016; Lee et al., 2001). Msps functions as a microtubule polymerase by binding to microtubule plus-ends (Lee et al., 2001). However, very little is known about their function in non-dividing cells. In this study, we demonstrate that microtubules in the primary protrusion of qNSCs are predominantly acentrosomal and oriented plus-end-out, distal to the cell body. We have identified Msps as a key regulator of NSC reactivation and microtubule dynamics. We show that E-cadherin (E-cad) is localized to the NSC-neuropil contact sites, in a Msps-dependent manner. E-cad is also intrinsically required for NSC reactivation. Our study, for the first time, has discovered microtubule plus-end-out orientation in the primary protrusion of qNSCs and a novel mechanism by which Msps-dependent acentrosomal microtubule assembly targets E-cad to NSC-niche contracts to promote NSC reactivation.

## RESULTS

### The centrosomes are immature and devoid of PCM proteins in quiescent NSCs from newly hatched larvae

We previously reported that microtubules marked by α-tubulin are present in the cellular extension of quiescent NSCs (qNSCs) (Li et al., 2017). How microtubules in qNSCs are nucleated is unknown. The centrosomes are a major microtubule organizing centre (MTOC) in most dividing cells including active NSCs and are composed of a pair of centrioles surrounded by pericentriolar material (PCM) proteins. To investigate whether functional centrosomes are responsible for the assembly of microtubules in qNSCs, we examined the localization of centrosomal proteins in wild-type qNSCs. Active NSCs in the central larval brain divide asymmetrically to generate differentiating daughter cells that eventually produce neurons towards the basal side, the inner layer of the larval central nervous system. Since the primary cellular extension of qNSCs in the central brain are connected with neuropils from neurons (Chell and Brand, 2010b; Truman and Bate, 1988a), we referred to the tip of the primary cellular extension as the basal side of qNSCs, while the opposite side of the cell, which is distal to primary protrusion and faces the surface of the larval brain as the apical region, and the site where the primary cellular extension is attached to the cell body as the neck region (illustrated in Fig 1A). First, we examined the localization of Asterless (Asl), a constitutive centriolar protein at the centrosomes in dividing cells (Varmark et al., 2007). Remarkably, at 2h ALH, Asl in qNSCs was observed mostly at the apical region distal to the cellular extension of qNSCs labelled by CD8-GFP driven by *grainy head* (*grh*)-Gal4 (Fig S1A; 81.8%, n=44). Occasionally, Asl-positive centrioles were observed at lateral region of the cell body (Fig S1A, 13.6%, n=44) or at the neck region (Fig S1A, 4.5%, n=44) where the primary protrusion was attached to the cell body in qNSCs. At 6h ALH and 16h ALH, we observed almost identical localization pattern of Asl (data not shown), suggesting that the centrosomes in wild-type qNSCs might be relatively stationary close to the apical surface in early larval stages. These data suggested that the centrosomes in qNSCs is positioned predominantly at apical region, which is distal to the primary cellular extension. Moreover, we found that the majority of wild-type qNSCs contained two Asl -positive centrioles (Fig S1A; 75%, n=44), while the rest of 25% (n=44) of qNSCs contained a single Asl - positive centriole. As centrioles are duplicated in S phase, qNSCs with two Asl-positive centrioles are presumably arrested in G2 phase, while qNSCs with a single Asl-positive centriole are likely arrested in G0 stage. This observation in the central brain is very similar to a recent report on the ratio of G2- and G0-arrested qNSCs in the VNC (Chell and Brand, 2010b).

**Figure 1.**
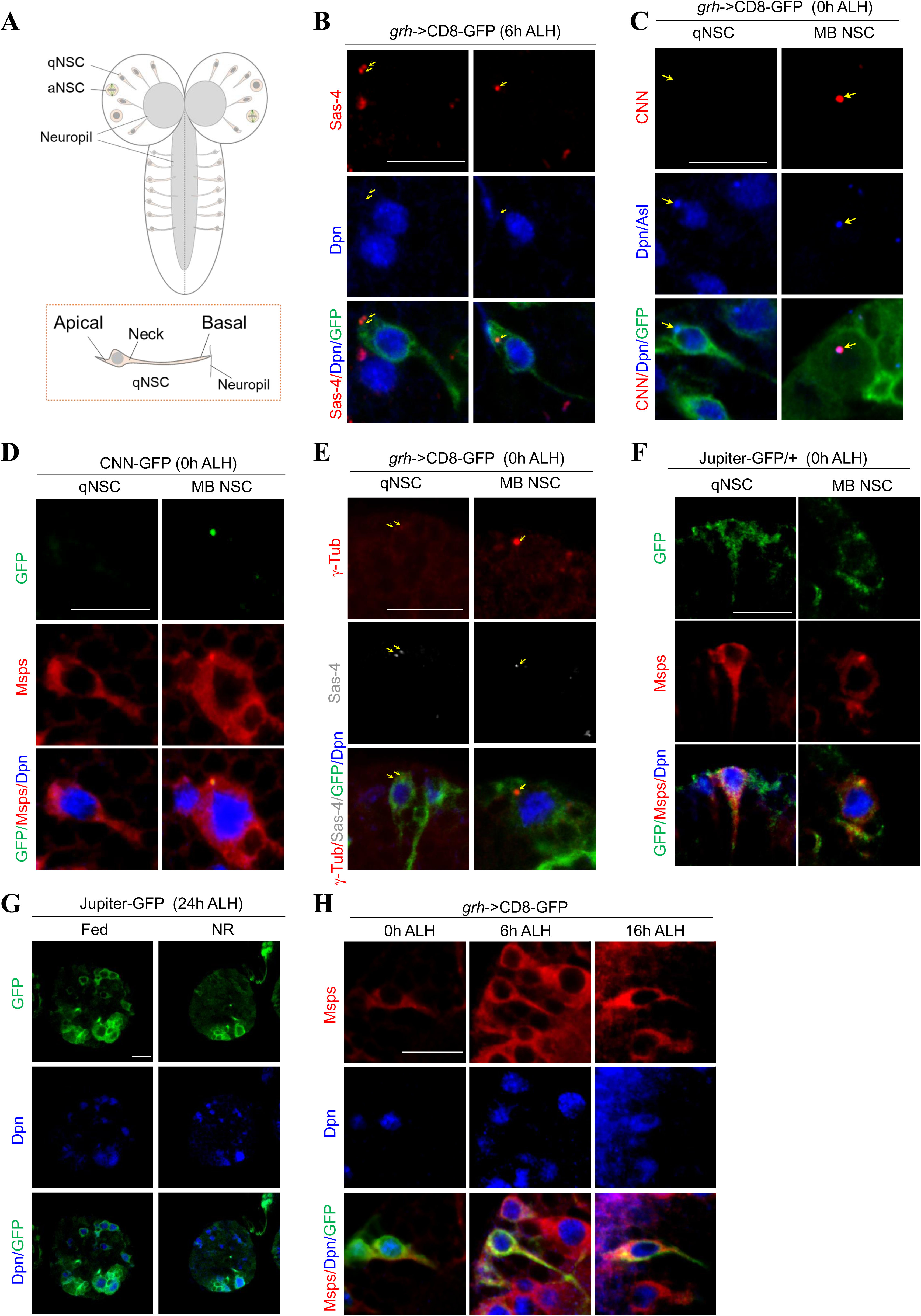
The centrosome in quiescent NSCs is immature. A) An illustration of the larval brain at early larval stages and the alignment of quiescent NSC along apico-basal axis. B) Larval brains at 6h ALH from *grh*-Gal4; UAS-CD8-GFP were labelled with Sas-4, Dpn and GFP. C) Larval brains at 0h ALH from *grh*-Gal4; UAS-CD8-GFP were labelled with Centrosomin (CNN), Asl, Dpn, and GFP. D) Wild-type larval brains expressing CNN-GFP (MiMIC line; BDSC#60266) at 0h ALH were labelled with GFP, Dpn, and Msps. A representative qNSC and interphase Mushroom body (MB) NSC were shown. E) Larval brains at 0h ALH from *grh*-Gal4; UAS-CD8-GFP were labelled with γ-tubulin (γ-tub), Sas-4, Dpn, and GFP. F) Larval brains at 0h ALH from Jupiter-GFP (G147) were labelled with GFP, Msps, and Dpn. G) Larval brains at 24h ALH from Jupiter-GFP were raised on normal food (fed) and food depleted of amino acids (nutritional restriction/NR) and labelled with GFP and Dpn. H) Larval brains at various time points from *grh*-Gal4; UAS-CD8-GFP were labelled with Msps, Dpn, and GFP. Quiescent NSCs at the central brain (CB) were shown (B-H). Arrows, centriole(s)/centrosomes (B-F). Scale bars: 10 µm.

Next, we examined another centriolar protein Sas-4 in qNSCs with a primary cellular protrusion labelled by CD8-GFP driven by *grh*-Gal4 at 6h ALH (Fig 1B; 100%, n=30). Very similar to Asl, vast majority of Sas-4 (Fig 1B; 83.3% n=30) was located at the apical region and distal to the primary protrusion of wild-type qNSCs, while 13.3% (n=30) were observed at lateral region of cell body. Only 3.3% of qNSCs (n=30) contained Sas-4-positive centriole(s) at the neck region of the primary protrusion, suggesting that the centrosomes may be occasionally motile within the cell body. Therefore, Sas-4 is primarily localized to the apical region, distal to the primary cellular extension in wild-type qNSCs. Similar to Asl, in wild-type qNSCs the majority of qNSCs contained two Sas-4 -positive centrioles (73.3%, n=30), while minority of qNSCs contain only a single Sas-4 - positive centriole. Taken together, we conclude that the centriolar proteins are predominantly localized to the apical region of wild-type qNSCs, distal to their primary cellular extension.

In dividing NSCs the centrosomes are mature during G2 phase and are responsible for the assembly of interphase microtubule aster (Rebollo et al., 2007; Rusan and Peifer, 2007). Given that the majority of qNSCs are arrested in G2 phase, we wondered whether the centrosomes in qNSCs were mature and had microtubule nucleation activity. To this end, we investigated whether PCM proteins were recruited to the centrosomes in qNSCs. The Pericentrin homologue named centrosomin (CNN) is an essential component of PCM and is required for centrosome assembly in dividing cells including active NSCs (Conduit and Raff, 2010). To ensure that NSCs we examined were definitely in quiescence, but not in a transition to reactivation from slightly older larvae, we examined CNN localization in qNSCs from newly hatched larvae (at 0h ALH). Surprisingly, CNN was completely absent in qNSCs at 0h ALH larval brains (Fig 1C left panels; 100%, n=53). By contrast, at 0h ALH in mushroom body (MB) NSCs that never enter quiescence, strong CNN signal was readily observed at the centrosomes (Fig 1C right panels; 100%, n=50). Likewise, CNN-GFP, in which endogenous CNN was tagged by GFP in a *Minos* mediated integration cassette (MiMIC) transposon insertion line (Venken et al., 2011), was also absent in qNSCs at 0h ALH (100%, n=29), but strongly expressed in MB NSCs at the same time point (Fig 1D; 100%, n=32). At 6h ALH CNN became weakly detectable in those qNSCs retaining primary cellular protrusion but with increased cell size (Fig S1B-C; 33.3%, n=42, average diameter 5.4 ± 0.2 μm), while the majority of qNSCs were still negative for CNN (69.7%, n=42, average diameter 3.8 ± 2.0 μm). These observations suggest that although subsequent recruitment of CNN to the centrosome is likely associated with the onset of NSC reactivation, the centrosomes in qNSCs at 0h ALH are likely immature and lack microtubule nucleation activity.

γ-tubulin (γ-tub) is a major microtubule nucleator in dividing cells such as cycling NSCs and non-dividing cells such as neurons. In dividing cells, γ-tub is a component of γ-tubulin ring complex (γ-TURC) robustly localized at the centrosomes. We wondered whether γ-tub was recruited to the centrosomes of qNSCs. We did a double labelling of γ-Tub and Sas-4, and found the γ-Tub could barely be detected in 89.1% (n=55) of qNSCs at 0h ALH, seemingly colocalizing with Sas-4 at the apical region, while γ-Tub was strongly localized to the centrosome in MB NSCs at the same time point (Fig 1E). γ-Tub was undetectable in the remaining of 10.9% (n=55) of qNSCs. γ-Tub signals were increased in intensity at the centrosome at 6h ALH but remained at low levels (Fig S1D, n=36). The nearly absent signal of γ-Tub in qNSCs at 0h ALH also supported the notion that the centrosome is not the major MTOC that is responsible for the microtubule assembly in the primary cellular protrusion of qNSCs.

To further investigate whether the centrosome is the major MTOC in qNSCs, we examined whether a microtubule aster could be detected in qNSCs. In dividing NSCs, an interphase microtubule aster is organized by the centrosome and could be detected by α-tubulin or β-tubulin. However, in qNSCs at 0h ALH, β-tubulin-Venus could be detected, but without forming a microtubule aster at the apical region where the centrosome is located (Fig S1E; 100%, n=45 in CB). Similarly, at both 6h- and 16h ALH, no microtubule aster at the apical region could be observed by α-tubulin (Fig S1F; 100%, n=37 in CB at 6h ALH; 100%, n=21 in CB at 16h ALH). Next, we examined Jupiter-GFP (G147), a protein trap line which labels microtubules and MTOC in dividing NSCs, was expressed under the control of endogenous *jupiter* promotor (Morin et al., 2001). At 0h ALH, Jupiter-GFP (G147) detected throughout the cytoplasm of qNSCs including the primary cellular protrusion, did not form any microtubule aster (Fig 1F; 100%, n=25 in CB). By contrast, microtubule aster marked by Jupiter-GFP was clearly observed in MB NSCs at 0h ALH (Fig 1f; 100%, n=40). Taken together, these observations indicate that the centrosomes in qNSCs from newly hatched larvae are immature and devoid of PCM proteins. Therefore, acentrosomal microtubule growth, rather than centrosomal microtubule growth, likely plays a major role in microtubule assembly in the cellular extension of qNSCs.

Next, we wondered whether microtubule polymerization in qNSCs is nutrition-dependent. We used Jupiter-GFP to label microtubules and raised the larvae at food depleted with amino acids (nutritional restriction/NR) for 24 hours. Strikingly, Jupiter-GFP signal intensity in the entire brain including in all Dpn-positive NSCs except for a few cells large in cell size that were presumably MB NSCs, was strongly reduced upon nutritional restriction compared with larval brains raised on fed condition (Fig 1G; 100%, NR, n=25 brain lobes [BL]; fed control, n=28 BL). This observation suggested that microtubule growth in qNSCs could be enhanced in the presence of nutrition.

Mini spindles (Msps) is a XMAP215/ch-TOG family protein and a key microtubule polymerase that promotes microtubule growth in dividing NSCs (Chen et al., 2016; Lee et al., 2001). However, the function of Msps in regulating microtubule assembly is poorly understood in non-dividing cells. To test whether Msps was expressed in the qNSCs, we examined the localization of Msps in qNSCs at 0h ALH. In qNSCs with the primary cellular protrusion labelled by CD8-GFP under the control of *grh*-Gal4, we detected cytoplasmic distribution of Msps throughout the qNSCs including the primary cellular protrusion (Fig 1H; 100%, n=36). This Msps signal observed in the wild-type qNSCs was specific, as Msps was undetectable in *msps^924^* and *msps^P18^* mutants at 0h ALH (Fig S3D). The distribution pattern of Msps in qNSCs at 16h ALH was similar to that in 0h ALH, but with overall higher levels (Fig 1H). These observations suggest that Msps is expressed in qNSCs from newly hatched larvae.

### Microtubules in the primary protrusion of qNSCs are predominantly orientated plus-end-out

Vertebrate neurons extend neurites with distinct microtubule orientation; Axons have a uniform arrangement of plus-end-out microtubules and dendrites have equal numbers of plus- and minus-end-out microtubules. In *Drosophila* neurons, axons have similar plus-end-out microtubule orientation, but microtubules in dendrites are primarily minus-end-out (Stone et al., 2008). We sought out to investigate microtubule orientation in the primary protrusion of qNSCs. First, we took advantage of two well-established microtubule polarity markers in *Drosophila* tissues including oocyte, epithelium, neuron and muscle (Clark et al., 1997). Kin-β-Galactosidase (Kin-β-Gal) marks plus-end microtubules and Nod-β-Gal marks minus-end microtubules by fusing the coiled-coil domain of a plus-end kinesin 1 motor protein or the motor domain of a minus-end kinesin-like Nod to β-gal (Clark et al., 1997). Since Kin-β-Gal was lethal when driven by *insc*-Gal4, we used *tubulin*-Gal80^ts^ to control the expression of Kin-β-Gal and Msps was used to label the primary protrusion of qNSCs. The embryos were kept at 18°C until larval hatching following by a shift to 29°C for 16 hours. In some qNSCs, Kin-β-Gal was undetectable (Fig 2A; 57%, n=77), likely due to its low expression level. In the remaining Dpn-positive qNSCs that Kin-β-Gal could be observed, it was localized mostly at the tip (Fig 2A; 72.7%, n=33) or in the middle of the primary cellular extension (27.3%, n=33) of qNSCs in the VNC. However, Kin-β-Gal was never observed at the apical region of qNSCs. Although at 0h ALH, Kin-β-Gal was mostly absent in qNSCs, presumably due to low level of expression, in qNSCs that it could be detected Kin-β-Gal was localized at the tip or in the medial region of the primary protrusion (Fig S2A; 90.6% within primary protrusion, 9.4% in the cell body, n=32), suggesting that microtubule orientation remains plus-end-out in qNSCs at different time points.

**Figure 2.**
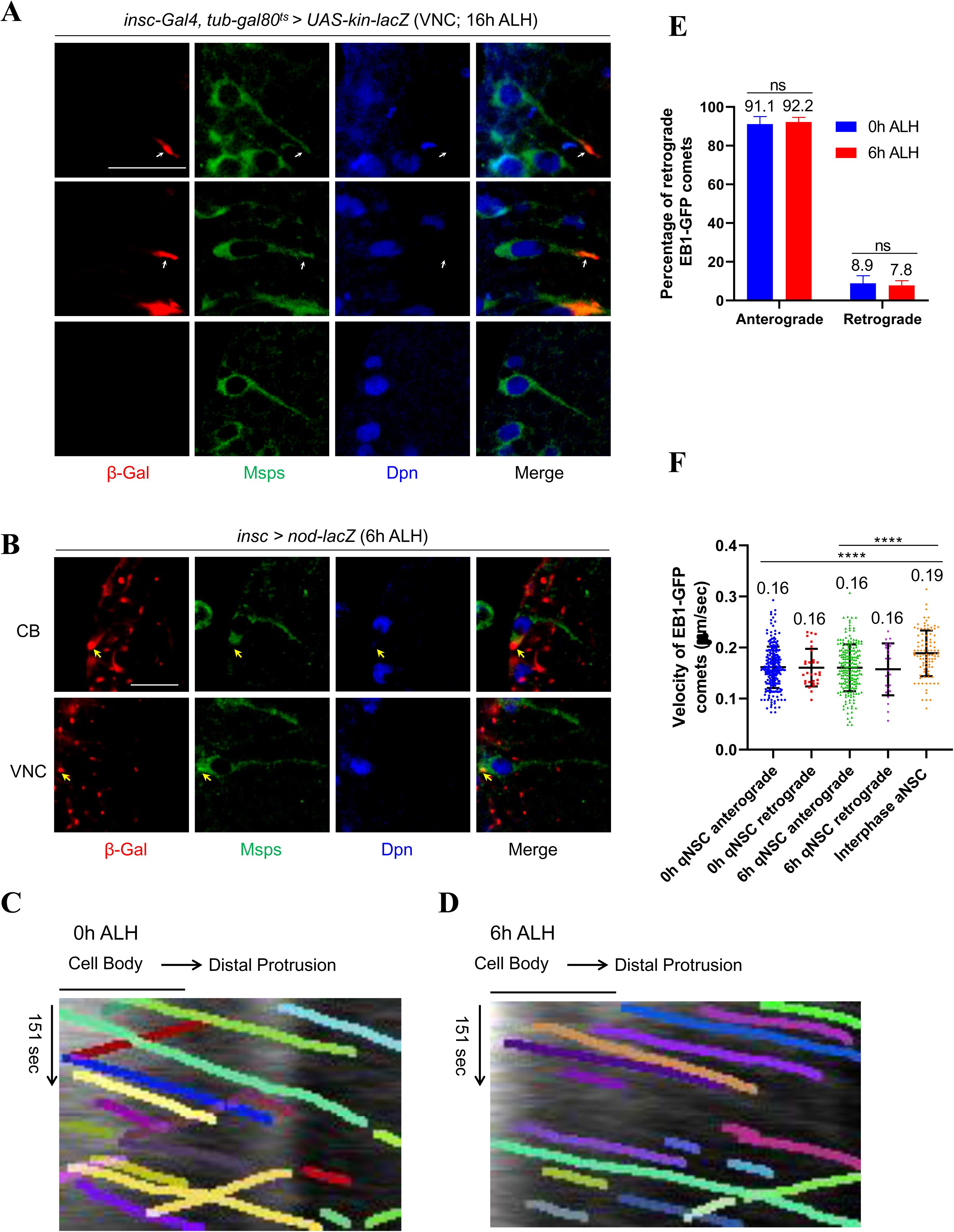
Microtubules in the cellular extension of qNSCs are predominantly plus-end-out orientated. A) Larval brains at 16h ALH, in which *kin-lacZ* was expressed under the control of *insc-*Gal4*, tub-*Gal80^ts^, were labelled with β-Gal, Msps, and Dpn. The primary protrusion of qNSCs were marked by Msps. Quiescent NSCs at the ventral nerve cord (VNC) were shown. B) Larval brains at 6h ALH from *insc*-Gal4; UAS-nod-lacZ were labelled with β-gal, Msps, and Dpn. Quiescent NSCs at both CB and VNC were shown. C) A representative kymograph depicting movements of EB1-GFP comets in the primary protrusion of qNSCs at 0h ALH. D) A representative kymograph tracking movements of EB1-GFP comets in the primary protrusion of qNSCs at 6h ALH. In c-d, the horizontal arrow indicates anterograde movement direction from cell body to the tip of the primary protrusion in qNSCs. E) Quantification graph of percentage of anterograde and retrograde movements of EB1-GFP comets in the primary protrusion of qNSCs at 0h ALH and 6h ALH. p=0.9096 (ns). F) Quantification graph of velocity of EB1-GFP comet movement in the protrusion of qNSCs at 0h ALH and 6h ALH, as well as in interphase NSCs (aNSC) at 72h ALH. ****p<0.0001. Scale bars: 10 µm.

Next, we examined the microtubule minus-end marker Nod-β-Gal in qNSCs with primary cellular extension marked by Msps. Remarkably, in both central brain (CB) and VNC at 6h ALH (Fig 2B; CB: 100%, n=42; VNC: 100%, n=37), Nod-β-Gal was predominantly localized at the apical region of qNSCs, co-localizing with centrioles marked by Asl (Fig S2C; 100%, n=28). At 0h ALH, the localization of Nod-β-Gal was predominantly localized at the apical region of qNSCs in both central brain (Fig S2B; apical, 94.3%; Neck region, 5.7%, n=35) and VNC (Fig S2B; Apical, 100%, n=25). Likewise, at 16h ALH, Nod-β-Gal was predominantly observed at the apical region of qNSCs (Fig S2D, 100%, n=24). Therefore, Kin-β-Gal is distributed at the basal side towards the tip of the primary protrusion, while Nod-β-Gal is at the apical side of qNSCs away from the primary protrusion, suggesting that microtubules in cellular extension of qNSCs are mostly oriented with plus-end-out distal to the cell body but towards the tip of the cellular extension.

To confirm the plus-end-out microtubule orientation of cellular protrusion of qNSCs, we proceeded to analyze End binding 1 (EB1), a plus-end tracking protein (+TIP) that binds to microtubule plus ends during microtubule growth (Vaughan, 2005). EB1-GFP was expressed under the control of *grh*-Gal4 to drive NSC specific expression at 6h ALH and dynamics of EB1-GFP comets were captured by live imaging followed by analysis of kymograph generated by imageJ in KymoButler (Jakobs et al., 2019). We termed the movement of EB1-GFP comets from the soma towards the basal side of qNSC as anterograde movement, and the opposite direction of movement as retrograde movement. Remarkably, at 0h ALH, 91.1% (n=28) of EB1-GFP comets in wild-type qNSCs displayed anterograde movement, while the remaining 8.9% (n=28) were retrograde movement (Fig 2C-E, Movie S2). The speed of anterograde and retrograde movements of EB1-GFP comets was essentially the same (Fig 2F; average velocity both at 0.16 μm/sec). These observations indicate that microtubules in the cellular extension of qNSCs are predominantly oriented plus-end-out distal to the cell body.

We wondered whether microtubule growth in the primary protrusion of qNSCs at a slightly later stage remained the same as that at 0h ALH. At 6h ALH, 92.2% (n=64) of EB1-GFP comets in wild-type qNSCs displayed anterograde movement, while the remaining 7.8% (n=64) were retrograde movement (Fig 2D-E; Movie S1). This suggested that the predominant microtubule orientation in the primary protrusion of qNSCs at 6h ALH is still plus-end-out, similar to that of 0h ALH. The speed between anterograde and retrograde movements of EB1-GFP comets at 6h ALH was indistinguishable (Fig 2F; Ave. velocity both at 0.16 μm/sec). The velocity of EB1-GFP comets in the primary protrusion of qNSCs at both 0h ALH and 6h ALH are slightly slower but very close to that of actively dividing NSCs (Fig 2F; 0.19 µm/sec in active NSCs; (Chen et al., 2016)). Taken together, microtubules in the primary protrusion of qNSCs are predominantly plus-end-out towards the tip of the protrusion and microtubule assembly in the primary protrusion of qNSCs is robust.

Because axons at presynaptic terminals oriented their microtubules plus-end-out, we wondered whether the primary protrusion of qNSCs resembles the structure of axons and expresses synaptic markers. nc82 antibody (anti-BRP) (Wagh et al., 2006), a widely used presynaptic marker, and Synaptotagmin (Syt), a synaptic vesicle-specific integral membrane protein found at the synaptic contact sites (Littleton et al., 1993) were both observed at neuropil, but absent in the primary protrusion of qNSC (Fig S2E; 100%, n=27 and Fig S2F; 100%, n=32). These observations suggest that despite similar plus-end-out microtubule orientation, the primary protrusion of qNSCs is functionally distinct from axons at the presynaptic terminals.

### Msps is critical for the reactivation of NSCs

Given that microtubule growth is unexpectedly robust in the primary protrusion of qNSCs, we tested the potential function of known microtubule regulators of dividing NSCs. We first tested ADP ribosylation factor like-2 (Arl2), a master regulator of microtubule assembly in dividing NSCs (Chen et al., 2016). We fed the larvae with EdU-containing food for 4h so that all cycling NSCs were incorporated with EdU as described previously (Li et al., 2017). Surprisingly, at 24h ALH, *arl2* knockdown (VDRC110627) driven by *insc*-Gal4 had no obvious defects in NSC reactivation (Fig S3A-B; EdU-negative NSCs in control, 3.9% of NSCs, n=15 BL; in *arl2* RNAi, 5.2%, n=11 BL). Overexpression of *arl2^T30N^*, a dominant-negative form of *arl2*, driven by *insc*-Gal4, only caused a very mild increase of EdU-negative NSCs from 3.9% (Fig S3A-B, control, n=15 BL) to 8.5% (*arl2^T30N^*, n=11 BL). In addition, depletion of centrosomes by *ana2* strong hypomophic alleles (Wang et al., 2011) also had no obvious impact to NSC reactivation (data not shown). Therefore, Arl2 or Ana2 are non-essential for NSC reactivation. Probably Arl2- and Ana2-dependent centrosomal microtubule growth is non-essential for NSC reactivation.

Because Msps is a microtubule polymerase that is expressed in qNSCs (Fig 1H), we investigated whether Msps is required for NSC reactivation. At 24h ALH, the vast majority of wild-type NSCs were reactivated and incorporated with EdU, while only 6.2% of NSCs were quiescent and negative for EdU (Fig 3A-B; n=13 BL). By contrast, the percentage of qNSCs that were EdU-negative was dramatically increased to 45.8% upon *msps* knockdown driven by *insc*-Gal4 (Fig 3A-B; n=13 BL), suggesting a significant delay of NSC reactivation. Next, we examined EdU incorporation of four loss-of-function alleles of *msps* including a protein null allele *msps^810^*, a strong hypomorphic allele *msps^924^* (Fengwei Yu, unpublished data) as well as two previously-reported hypomorphic *msps* alleles (Chen et al., 2016; Cullen et al., 1999). Remarkably, at 24h ALH, 86.2% (Fig 3C-D; n=13 BL) of NSCs in *msps^810^* failed to incorporate EdU, compared with only 12.5% (Fig 3C-D; n=13 BL) of NSCs without EdU incorporation in wild-type. This observation suggested that most of NSCs in *msps^810^* failed to exit quiescence. Similarly, all three other alleles of *msps* also displayed prominent NSC reactivation phenotype (Fig 3C-D; 59.2%, n=17 BL in *msps^924^*; 46.9%, n=14 BL in *msps*^P18^; and 64.7%, n=6 BL in *msps*^P^). To confirm the reactivation defects, we measured the cell diameter at 24h ALH, as qNSCs have a cell diameter of ∼4 μm, while reactivating NSCs undergo their first cell division when they reach the cell diameter of ∼7 μm (Chell and Brand, 2010a). At 24h ALH, the average cell diameter in *msps^924^* NSCs and *msps^P^* were 4.9 μm and 4.7 μm respectively, significantly smaller than 7.6 μm in wild-type NSCs (Fig 3C,E). Since Msps is required for Mira localization at cortex in NSCs (Chen et al., 2016) and Mira in qNSCs with *msps* depletion was cytoplasmic and could no longer label the primary protrusion clearly, we used CD8-GFP controlled by *grh*-Gal4 to mark the primary protrusion in *msps* mutants. At 24h ALH, there were more qNSCs with primary protrusion observed upon *msps* depletion (Fig 3F; *msps^924^*: 24.5%, n=6 BL; *msps*^P18^: 29.0%, n=4 BL), compared with the control (4.3%, n=5 BL). The NSC reactivation phenotype observed in *msps^924^* mutants was unlikely primarily due to a failure in mitosis, as even at 24h ALH, ectopic NSCs were observed in *msps^924^*, presumably due to symmetric division of *msps^924^*, a phenotype described for *msps^P18^* at late larval stages (Chen et al., 2016). Msps was undetectable in *msps^924^* and *msps^P18^* NSCs at 24h ALH and strongly reduced upon *msps* RNAi knockdown (Fig S3C), suggesting that Msps was sufficiently depleted at these conditions. Moreover, the EdU incorporation defects in *msps^810^* was nearly fully restored by the expression of a wild-type genomic *msps* (Fig 3C-D; n=12 BL). These observations indicate that Msps, but not Arl2, is essential for NSC reactivation and that Msps functions intrinsically in NSCs to promote their reactivation.

**Figure 3.**
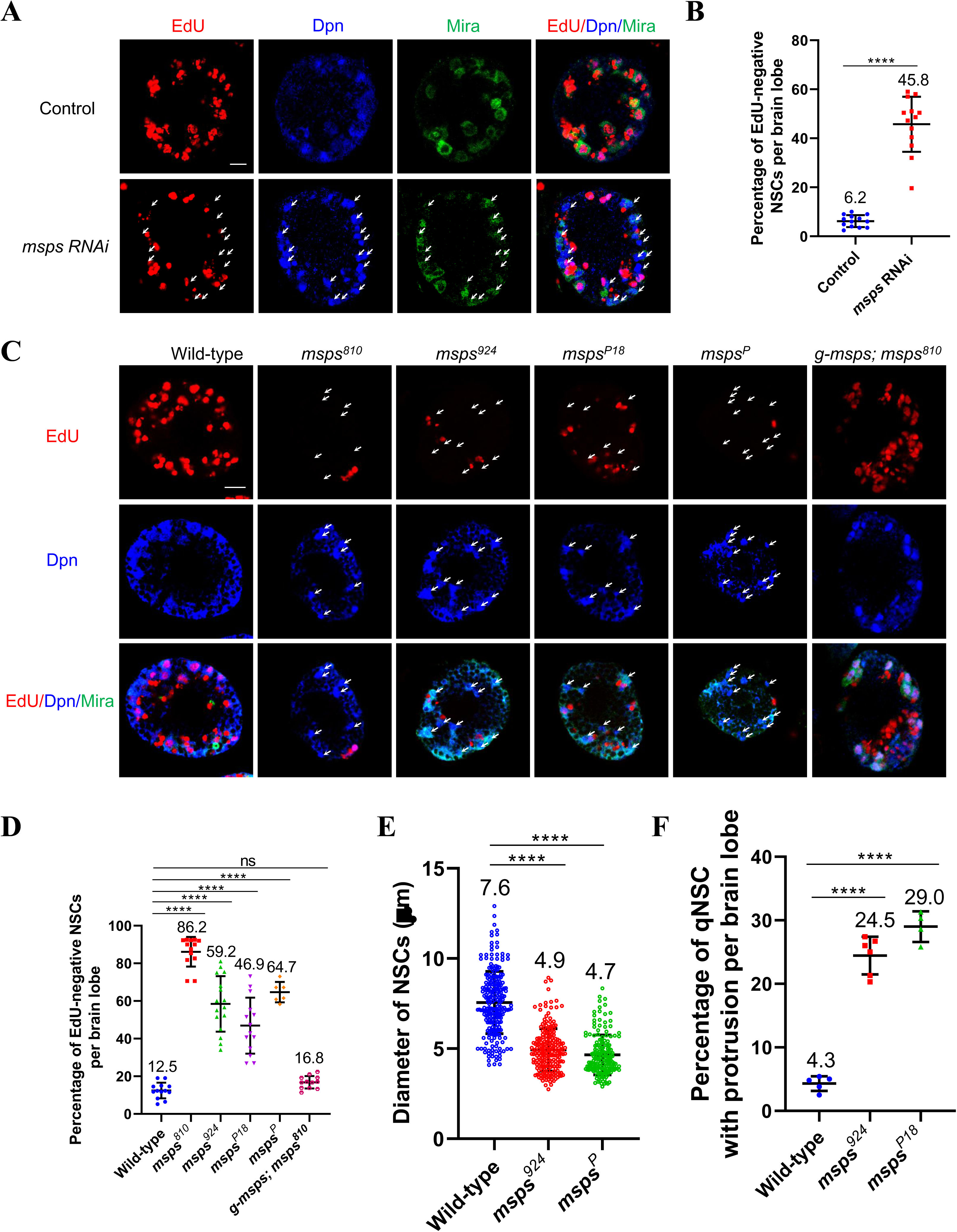
Msps is essential for NSC reactivation. A) Larval brains at 24h ALH from control (*insc*-Gal4; *UAS-dicer2*/*UAS-β-Gal* RNAi) and *msps* RNAi (VDRC#21982) controlled under *insc*-Gal4 were analyzed for EdU incorporation. NSCs were marked by Dpn and Mira. B) Quantification graph of EdU-negative NSCs per brain lobe for genotypes in (a). ****p< 0.0001. C) Larval brains at 24h ALH from wild-type and various *msps* loss-of-function alleles and *msps^810^* with a genomic rescue construct (*g-msps*) were analyzed for EdU incorporation. NSCs were marked by Dpn and Mira., D) Quantification graph of EdU-negative NSCs per brain lobe for genotypes in (c). ****p< 0.0001; p=0.7502 (ns). F) Quantification graph of diameter of the cell body in NSCs at 24h ALH from various genotypes. ****p< 0.0001. F) Quantification graph of percentage of qNSCs with primary protrusion in wild-type, *msps^924^*, and *msps^P18^.* The protrusion was labelled by *grh>CD8-GFP*. Scale bars: 10 µm.

### Msps is critical for acentrosomal microtubule assembly in the primary protrusion of qNSCs

We investigated whether Msps was important for microtubule growth in the primary protrusion of qNSCs. First, we analysed microtubule growth by tracking the movement of EB1-GFP comets in the primary protrusion of qNSCs upon *msps* depletion. At 6h ALH, EB1-GFP comets were almost completely lost in the primary protrusion of qNSCs from a hypomorphic allele *msps^P18^* and a trans-heterozygous mutant *msps^P18/P^* (Fig 4A-B, Movies S3-S4; *msps^P18^*, 0.05 fold, n=18; *msps^P18/P^*, 0.02 fold, n=21, compared with control, 1, n=26). This result indicates that Msps is critical for microtubule polymerization in the primary protrusion of qNSCs. Next, we examined Nod-β-Gal localization in qNSCs upon *msps* RNAi knockdown. In control qNSCs labelled by Dpn and *grh*-Gal4 UAS-CD8-GFP, Nod-β-Gal was concentrated at the apical region of the primary protrusion (Fig 4C-D, n=60). By contrast, in 85.1% of qNSCs upon *msps* RNAi knockdown, Nod-β-Gal was delocalized from the apical region and distributed around the cell body, even observed at the primary protrusion (Fig 4C-D, n=68). This striking phenotype suggested that the microtubule orientation might be altered in qNSCs upon *msps* depletion.

**Figure 4.**
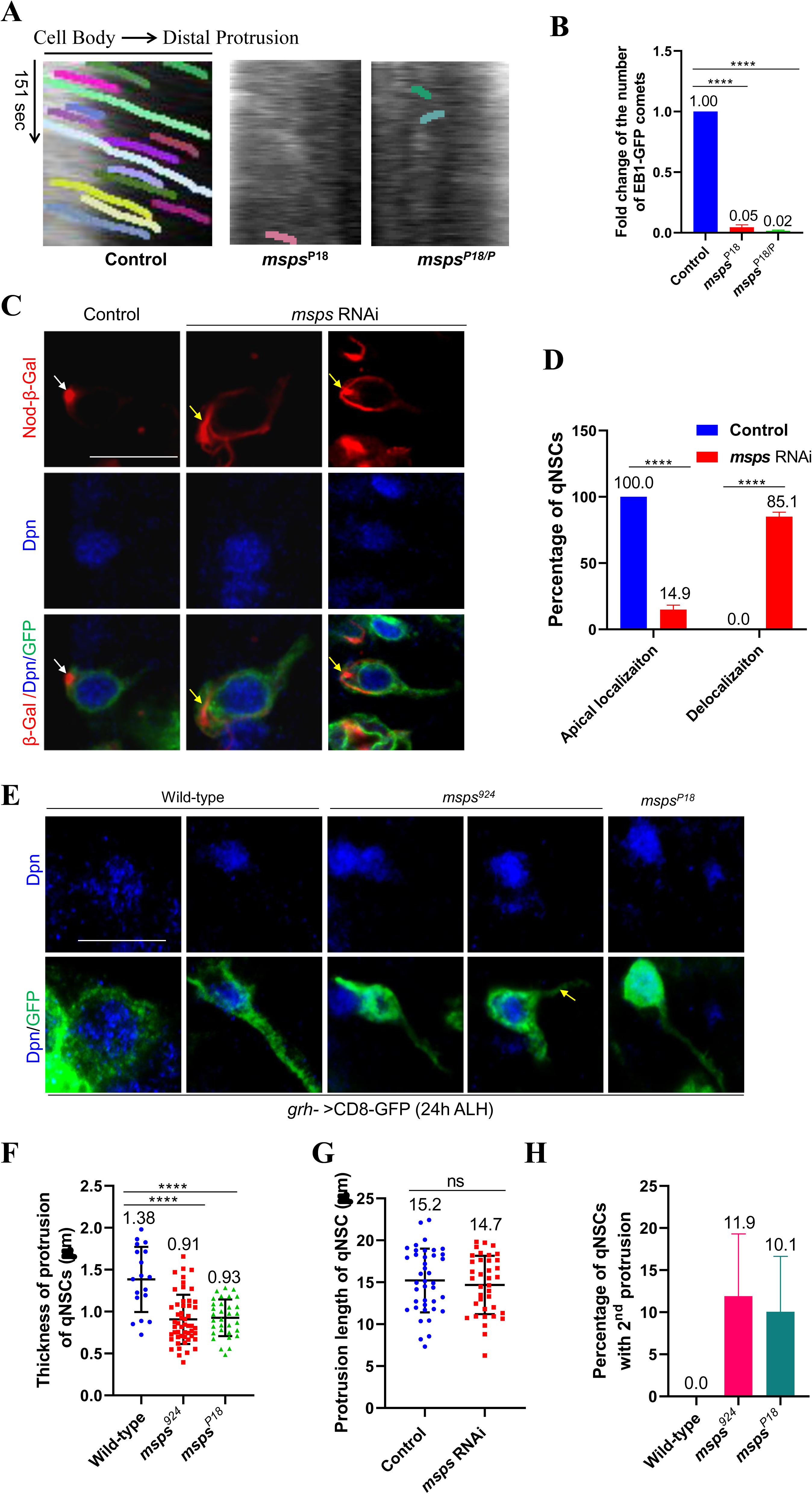
Msps regulates microtubule assembly in the primary protrusion of qNSCs. A) Kymograph of EB-GFP comets movement in the primary protrusion of qNSCs from control, *msps^P18^* and *msps^P18/P^* with EB1-GFP expressed under *grh*-Gal4 at 6h ALH, the horizontal arrow indicates anterograde movement direction from cell body to the tip of the primary protrusion in qNSCs. B) Quantification graph of fold changes of number of EB1-GFP comets in the primary protrusion of qNSCs 6h ALH from various genotypes compared with control in (a). ****p<0.0001.C) Larval brains at 16h ALH from control (*grh*-Gal4 *UAS-CD8-GFP*; *UAS-dicer2* + *UAS*-*nod-lacZ*) and *msps* RNAi, *UAS-nod-lacZ* (VDRC#21982) with *grh*-Gal4 *UAS-CD8-GFP*; *UAS*-*dicer2* were labelled with β-Gal, Dpn, and GFP. Quiescent NSCs at the central brain (CB) were shown. D) Quantification graph of Nod-β-Gal localization in qNSCs from genotypes in (c). ****p<0.0001.E) Larval brains at 24h ALH from wild-type, *msps^924^*, and *msps^P18^* expressing *grh>*CD8-GFP were labelled with Dpn and GFP. F) Quantification graph of thickness of the primary protrusion of qNSCs from wild-type, *msps^924^*, and *msps^P18^* expressing *grh>*CD8-GFP. The thickness was measured at the middle point of the primary protrusion. **** p<0.0001. G) Quantification graph of the length of primary protrusion in qNSCs in control (*β-Gal* RNAi) and *msps* RNAi (VDRC#21982) under the control of *grh*-Gal4 with *UAS-CD8-GFP UAS-dicer2*. P=0.5257 (ns). H) Quantification graph of qNSCs extended two major protrusions labelled by *grh>CD8-GFP* in wild-type, *msps^924^*, and *msps^P18^*. Scale bars (a, f): 10 µm.

Given that Msps is critically required for microtubule growth in the primary protrusion of qNSC, we examined whether *msps* depletion resulted in morphological defects in the primary protrusion of qNSCs. The thickness of the primary protrusion was measured at the middle position of the primary protrusion marked by *grh*>CD8-GFP. Loss-of-*msps* resulted in dramatic thinning of the primary protrusion in qNSCs (Fig 4E-F; control, 1.38 ± 0.39 μm, n=18; *msps^924^*, 0.91 ± 0.29 μm, n=52; *msps*^P18^, 0.93 ± 0.22 μm, n=32). However, the length of the primary protrusion in qNSCs (VNC) upon *msps* RNAi knockdown at 16h ALH was not statistically different from that of the control (Fig 4G; Control,15.2 ± 3.7 μm, n=40; *msps* RNAi, 14.7 ± 3.4 μm, n= 37), which is likely due to the distance between the cell body of qNSCs and neuropil is relatively constant. In wild-type, qNSCs occasionally extend a second protrusion toward the apical side or lateral side of the cell body in the first a few hours after larval hatching, but this structure is not seen in qNSCs at 24h ALH (Fig 4E,H; 0% secondary protrusion, n=18). However, in *msps* mutant qNSCs, the secondary protrusion was readily observed (Fig 4E,H; *msps^924^*, 11.9 ± 7.4%, n=65*; msps*^P18^, 10.1 ± 6.6%, n=71). More frequent extension of secondary protrusion seemed to associate with a weakened primary protrusion in *msps^924^* NSCs.

To further exclude the possibility that the centrosomes are potentially contributing to the microtubule assembly in the primary protrusion of qNSCs, we examined the EB1-GFP comets in qNSCs upon *sas4* or *ana2* depletion, which were known to result in defects of centrosome formation. Strikingly, loss of centriolar protein *sas-4* or *ana2* didn’t apparently disrupt MT assembly in the primary protrusion of qNSCs. At 6h ALH, the number of EB1-GFP comets in the primary protrusion of qNSCs were similar among control, *sas4* RNAi and *ana2* RNAi in the heterozygous *ana2^719^* background (Fig S4C-D, Movies S5-S6; control, 1, n=10; *sas-4* RNAi, 1.27 fold, n=10; *ana2* RNAi *ana2^719^/+*, 1.03 fold, n=16). Sas4 and Ana2 were mostly undetectable in qNSCs from *sas4* RNAi (94.4%, n=18) and *ana2* RNAi *ana2^719^/+* (86.3%, n=22) (Fig S4A-B), suggesting an efficient knockdown. However, depletion of *sas4* or *ana2* resulted in a very mild phenotype or no phenotype of NSC reactivation (Fig S4E-G). These observations indicate that the centrosomes are non-essential for microtubule assembly in the primary protrusion in qNSCs.

Taken together, Msps is critical for acentrosomal microtubule growth in the primary protrusion of qNSCs.

### E-cad localizes to NSC-neuropil contact points, in a Msps-dependent manner

Although the primary protrusion of qNSCs is known to reach neuropil (Chell and Brand, 2010b), proteins that localize to the NSC-neuropil contact points have not been identified. We examined the localization of E-cad/Shotgun in qNSCs, as E-cad is a cell adhesion molecule that has homophilic interactions and often localizes to cell-cell contacts. The outlines of qNSCs could be clearly labelled by CD8-GFP expressed under the control of *grh*-Gal4, where their basal tips join at the bulk of CD8-GFP-positive neuropil (Fig 5A). We found that in wild-type VNCs at 24h ALH, E-cad was observed at the cell cortex of qNSCs and formed an endfeet-like structure at the tip of the protrusion and at the surface of neuropil where qNSCs contact neuropil (Fig 5A; n=95). However, E-cad was only very weakly observed within the bulk body of neuropil (Fig 5A). E-cad localization at NSC-neuropil contact sites and within NSCs were dramatically reduced upon knocking down E-cad in NSCs (Fig S5B-C), suggesting that these localizations in wild-type qNSCs were specific for E-cad.

**Figure 5.**
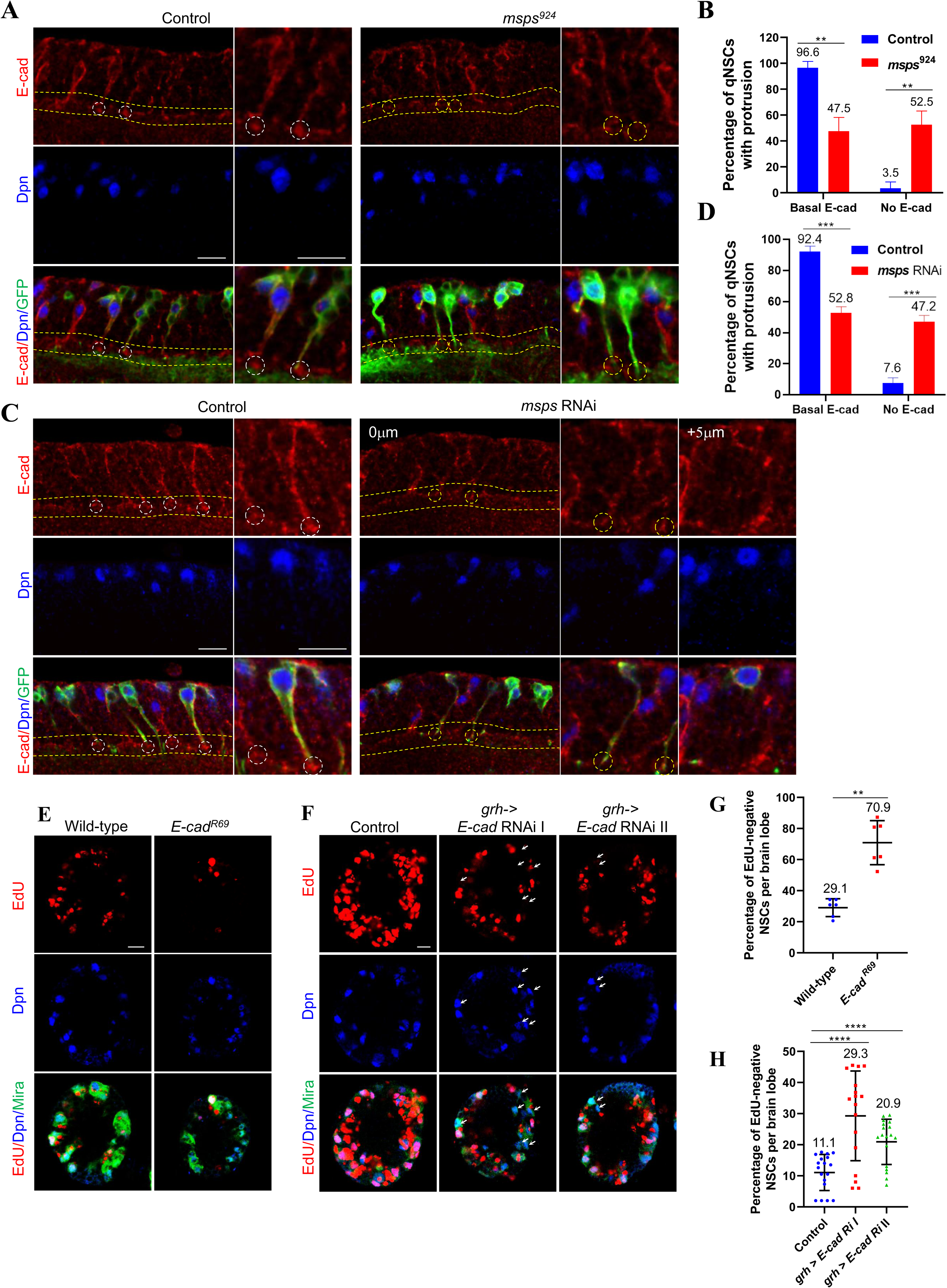
Delocalization of E-Cadherin at NSC-neuropil contact sites in qNSCs upon *msps* depletion. A) Larval VNCs at 24h ALH from control (*grh-*Gal4 *UAS-CD8-GFP*) and *msps^924^* with *grh-*Gal4 *UAS-CD8-GFP* were labelled with E-cadherin, Dpn, and GFP. B) Quantification of E-Cadherin basal localization at NSC-neuropil contact sites in qNSCs from genotypes in (A). “No E-cad” means absent or strongly reduced E-cad observed at basal region of qNSCs. **p=0.0080. C) Larval VNCs at 16h ALH from control (*grh-*Gal4 *UAS-CD8-GFP; UAS-Dicer2 + UAS-β-Gal* RNAi) and *msps* RNAi with *grh-*Gal4 *UAS-CD8-GFP; UAS-Dicer2* were labelled with E-cadherin, Dpn, and GFP. The single-scanning image “+5 μm” showed cortical localization of E-Cad in the cell body of quiescent NSCs, suggesting that there is no gross reduction of E-cad in NSCs upon *msps* depletion. D) Quantification of E-Cadherin basal localization at NSC-neuropil contact sites in qNSCs from genotypes in (C). “No E-cad” means absent or strongly reduced E-cad observed at NSC-neuropil contact points. ***p=0.0008. E) Larval brains at 16h ALH from Wild-type and *E-cad^R69^* were analyzed for EdU incorporation and larval brains were labelled with EdU, Dpn and Mira. F) Larval brains at 24h ALH from control (*grh*-Gal4; *UAS-dicer2*/*UAS-β-Gal* RNAi), *E-cad* Ri I (UAS-*E-cad* RNAi (BDSC#32904) and *E-cad* Ri II (UAS-*E-cad* RNAi (BDSC#38207) controlled under *grh*-Gal4; *UAS-Dicer2* were analyzed for EdU incorporation. G) Quantification graph of EdU-negative NSCs per brain lobe for genotypes in (E). ****p<0.0001. H) Quantification graph of EdU-negative NSCs per brain lobe for genotypes in (F). **p<0.0022. Scale bars: 10 µm.

Next, we tested whether Msps is required for the E-cad localization at the NSC-neuropil contact sites. Strikingly, basal NSC-neuropil contact sites were lost in 52.5% of *msps^924^* qNSCs (Fig 5A-B; n=30) marked by *grh*>CD8-GFP, compared with the control qNSCs (Fig 5A-B; 92.4%, n=77). Likewise, E-cad was delocalized from NSC-neuropil contact sites in 47.2% (Fig 5C-D; n=44) of qNSCs upon *msps* RNAi knockdown, compared with 92.4% E-cad localization at the NSC-neuropil contact sites in control (Fig 5C-D; n=39). Therefore, E-cad localizes to NSC-neuropil contact sites, which requires functional Msps within NSCs.

E-cad was previously reported to act in glia cells to promote the proliferation of NSCs (Dumstrei et al., 2003). However, it was unknown whether E-Cad is required for NSC reactivation, as previous analyses on E-cad were carried out in 3^rd^ instar larval brains, when NSCs proliferation is independent of dietary amino acids. To test the role of E-cad in NSC reactivation, we examined *E-cad^R69^*, a known null allele resulted from a P-element excision that removes the translation start site of *E-cad* (Godt and Tepass, 1998), at 16h ALH, as the homozygotes did not survive to 24 h ALH. At 16h ALH, majority of NSCs from *E-cad^R69^* failed to incorporate EdU (Fig 5E,G; 70.9%, n=6 BL), compared with 29.1% EdU-negative NSCs in the wild-type control (Fig 5E,G; n=6 BL). In addition, 27.5% (Fig S5D; n=19) of *E-cad^R69^* NSCs still retained primary protrusion, compared with 15.4% (n=16) in wild-type. And this phenotype was completely restored by ectopic E-cad-GFP expression under the control of Ubi-p63E promoter (Fig S5D; 16.2%, n=9 in rescued animals). Moreover, at 16h ALH, there was a significant reduction of cell diameter in *E-cad^R69^* (Fig S5E; 5.1 ± 1.4 μm, n=131), compared with wild-type (6.4 ± 1.9 μm, n=110). Ectopic expression of E-cad-GFP restored the cell growth defect in *E-cad^R69^* NSCs (Fig S5E; 6.4 ±1.6 μm, n=114). Furthermore, mitotic index was significantly reduced in *E-cad^R69^* mutant brains at 16h ALH (Fig S5F; 9.4%, n=9 BL), compared with that of wild-type (21.0%, n=8 BL). Taken together, E-cad is required for NSC reactivation.

Given that E-cad is localized to NSC-neuropil contact sites, we investigated whether E-cad was required in NSCs to promote NSC reactivation. Upon knocking down of *E-cad* by two independent RNAi lines driven by *grh*-Gal4 at 24h ALH, there were a significant increase of EdU-negative NSCs (Fig 5F,H; *E-cad* RNAi I, 29.3%, n=17 BL; *E-cad* RNAi II, 20.9%, n=18 BL), compared with 11.1% EdU-negative NSCs in the control (Fig 5F,H; n=18 BL). Moreover, there were a significant increase of percentage of qNSCs retaining primary protrusion upon *E-cad* knockdown under *grh*-Gal4 (Fig S5G; *E-cad* RNAi I, 15.8%, n=9 BL; *E-cad* RNAi II, 11.7%, n=8 BL), compared with control (Fig 5G; 5.6%, n=9 BL). In addition, cell diameter of NSCs from *E-cad* RNAi knockdown under *grh*-Gal4 was significantly decreased (Fig S5H; *E*-*cad* RNAi I, 5.6 ± 1.6 μm, n=312; *E-cad* RNAi II, 6.1 ± 1.4 μm, n=353), compared with control (7.0 ± 1.4 μm, n=343). These results support the conclusion that E-cad acts in NSCs to promote their reactivation.

## DISCUSSION

In this study, we demonstrate that microtubules in the primary protrusion of qNSCs are predominantly acentrosomal and oriented plus-end-out, distal to the cell body. We have identified Msps/XMAP215 as a first key regulator of acentrosomal microtubule assembly in quiescent NSCs. We also reported the localization of E-cad at NSC-neuropil contact sites. Msps is important for the targeting of E-cad to NSC-niche contact points. Loss-of-function of *msps* and *E-Cad* in NSCs results in a failure of NSC reactivation. Our study, for the first time, demonstrate microtubule plus-end-out orientation in the primary protrusion of quiescent NSCs and a novel mechanism by which Msps governs NSC reactivation via targeting E-cad to NSC-neuropil contact sites.

Here we show that in the primary protrusion of wild-type qNSCs, the speed of EB1-GFP comets movement is slightly slower but very close to that in dividing NSCs, but dramatically faster than that reported in the dendrites of sensory neurons (∼0.1 µm/sec) (Wang et al., 2019). This result suggests that microtubule assembly in the protrusion is unexpectedly robust. Is this microtubule growth in qNSCs dependent on the centrosomes? The centrosomes are key MTOCs of dividing cells including active *Drosophila* NSCs. In cycling cells, PCM proteins are recruited to the centrosome(s) throughout the cell cycle with highest levels at the centrosomes during mitosis (Rebollo et al., 2009). In dividing larval brain NSCs, apical centrosome organizes a microtubule aster that stays at the apical cortex for most of the cell cycle, while the other centrosome loses PCM and microtubule-organizing activity and move dynamically until shortly before mitosis (Rebollo et al., 2007; Rusan and Peifer, 2007). We show that the centrosome in qNSCs have distinct behaviour from those in active NSCs. In newly hatched larvae, both centrosomes are mostly located at the apical region of qNSCs and are inactive devoid of PCM proteins such as CNN and γ-tub. Since these qNSCs have already extended their primary protrusion, the centrosomes are unlikely the MTOC responsible for organizing the primary protrusion. Consistent with these observations, microtubule aster could not be observed in qNSCs in early larval stages, suggesting the inability of microtubule nucleation by the centrosomes at this stage. Remarkably, tracking EB1-GFP comets in centrosome-deficit qNSCs indicates that microtubule growth in the primary protrusion of qNSCs is primarily independent of the centrosomes. Therefore, our study, for the first time, indicates that microtubule growth in the primary protrusion of qNSCs is surprisingly robust and mostly acentrosomal.

Microtubules are polarized and display uniform plus-end-out orientation in axons and minus-end-out orientation in dendrites in *Drosophila* neurons (Stone et al., 2008). We show that in the primary protrusion of qNSCs, microtubules are predominantly oriented with their plus-end-out (plus-end distal to the cell body). This is quite similar to the microtubule plus-end-out orientation in axons of both vertebrate and invertebrate neurons. Another similarity between the microtubule organization in the primary protrusion of qNSCs and axons is the inactive centrosomes in both cell types (Nguyen et al., 2011). However, we show that the centrosomes in wild-type qNSCs have a stereotypic position at the apical region in early larval stages, distinct from the centrosome of neuron without a consistent position (Nguyen et al., 2011). In addition, the primary protrusion of qNSCs project to and contact neuropil. Is the NSC-neuropil contact similar to synapses? The primary protrusion of qNSCs does not expression synaptic markers such as Synaptotagmin, suggesting a distinct composition from synapses.

We have identified Msps/MAP215 as a key regulator of acentrosomal microtubule growth in the primary protrusion of qNSCs. Msps/MAP215 directly binds to tubulin dimer via its tumor-overexpressed gene (TOG) domains to promote microtubule polymerization^35^. In dividing NSCs, Msps is mainly detected at the centrosomes in both interphase and mitosis and loss-of-*msps* led to the formation of a shorter spindle and NSC polarity defects (Chen et al., 2016). Unlike the absence of PCM proteins in newly hatched larvae, Msps was already seen in the cytoplasm of qNSCs including the primary protrusion. *msps* depletion resulted in a gross loss of EB1-GFP comets in the protrusion of qNSCs, an indication of loss of microtubule growth. Surprisingly, Arl2, which is essential for microtubule growth and centrosomal localization of Msps in dividing NSCs (Chen et al., 2016), appears to be not important for NSC reactivation. It is most likely that Msps regulates acentrosomal microtubule growth in qNSCs, independent of Arl2 that is more critical for centrosomal microtubule growth. Although the centrosome is immature in qNSCs from newly hatched larvae, the intensity of PCM at the centrosomes subsequently increases but remains low in reactivating NSCs. Perhaps the centrosome only becomes mature and functional following the retraction of the primary protrusion of NSCs. It is unknown whether the presence of the primary protrusion prevents centrosomes from maturation.

Microtubule plus-end-out orientation in axons of neurons is important for axonal growth and transport (Voelzmann et al., 2016). What is the function of plus-end-out polarity in the primary protrusion of qNSCs? We show that Msps is important for the formation of primary protrusion of qNSCs and its depletion results in dramatic thinning of the protrusion. Therefore, Msps-dependent microtubule growth provides a structural support to the formation of primary protrusion. Although the presence of primary protrusion is believed to be a hallmark of qNSCs, we provide evidence that Msps-dependent microtubule assembly in the protrusion is likely required for NSC reactivation. Primary protrusion of qNSCs directly contacts with neuropil, however, proteins at NSC-neuropil contact sites were previously unknown. Cell adhesion molecule E-cad, abundantly expressed in 3^rd^ instar larval NSCs, is often localized to cell-cell contacts and was recently reported to localize to MB NSC-cortex glia contact in the adult *Drosophila* brain (Doyle et al., 2017; Dumstrei et al., 2003). We show, for the first time, that E-cad forms an endfeet-like structure at NSC-neuropil contact sites. Moreover, Msps is important for targeting E-cad to the cell junctions where qNSCs contact their niche at the neuropil. E-cad was shown previously to be required in glia to promote NSC reactivation (Dumstrei et al., 2003). We further show that E-cad is intrinsically required for NSC reactivation.

Recently, microtubule-based nanotubes was shown to mediate signalling between *Drosophila* male germline stem cells and their niche (Inaba et al., 2015). The structure of the primary protrusion in qNSCs is distinct from that of nanotubes, as the latter lacks acetylated tubulin and is much thinner and shorter (∼0.4 μm in thickness and ∼3 μm in length) than the former (∼1.4 μm in thickness and ∼15 μm in length). Moreover, qNSC primary protrusion appears to be distinct from primary cilia, as the latter is assembled/attached from the basal body, which is derived from the mother centriole (Seeley and Nachury, 2010). The primary protrusion of qNSCs also differs from cytonemes and tunnelling nanotubes that were up to 700-1000 µm in length and mediated long-range signalling between cells (Roy et al., 2014; Rustom et al., 2004). We propose that the primary protrusion of qNSCs is a novel type of cellular protrusion that potentially mediates NSC-neuropil communication. Our finding that E-cad localizes to the NSC-neuropil contact sites and is required for NSC reactivation suggests that the neuropil contacted by the primary protrusion of quiescent NSCs likely functions as a new niche for promoting NSC reactivation.

Taken together, we propose that Msps is essential for acentrosomal microtubule growth in qNSCs, which facilitates targeting of E-cad to NSC-niche contact points to promote NSC reactivation. Our findings may be a general paradigm that could be applied to other types of quiescent stem cells in both *Drosophila* and mammalian systems.

## ACKNOWLEDGMENTS

We thank F. Yu, A. Brand, C. Gonzalez, J. Raff, E. Schejter, T. Megraw, T. Lee, J. Knoblich, F Matsuzaki, and Y. Jan, the Bloomington Drosophila Stock Center, Vienna Drosophila Resource Center, Kyoto stock centre DGGR, and the Developmental Studies Hybridoma Bank for fly stocks and antibodies. This work is supported by Ministry of Health-Singapore National Medical Research Council MOH-000143 (MOH-OFIRG18may-0004) to H.W.

## AUTHOR CONTRIBUTIONS

Conceptualization, H.W., Q.D. and F.Y.; Methodology, Data curation, and Formal analysis, Q.D, Y.S.T, and L.Y.C.; Writing-Original draft, Q.D. and H.W.; Writing-Review & Editing, H.W., Q.D, and F.Y; Funding Acquisition, H.W.; Resources, H.W. and F.Y; Supervision, H.W.

## METHODS

### Fly stocks and genetics

Fly stocks and genetic crosses were raised at 25°C unless otherwise stated. Fly stocks were kept in vials or bottles containing standard fly food (0.8% Drosophila agar, 5.8% Cornmeal, 5.1% Dextrose and 2.4% Brewer’s yeast). The following fly strains were used in this study: *insc-*Gal4 (BDSC#8751; 1407-Gal4), *grh-*Gal4 (A. Brand), *insc-*Gal4 *tub-*Gal80^ts^*, msps^924^* (F. Yu), *msps^810^* (F. Yu)*, msps^P18 (Chen et al., 2016)^, msps^P^* (Cullen et al., 1999), *g-msps* (HN267) (Cullen et al., 1999), *UAS-Kin-β-gal* (Clark et al., 1997), *UAS-arl2^T30N^*/*TM6B Tb* (Chen et al., 2016), *Jupiter-GFP* (G147), *UAS-β-tub-Venus/CyOβ,*, *UAS-GFP-msps/TM6B Tb* (F. Yu). The following stocks were obtained from Bloomington Drosophila Stock Center (BDSC): *UAS-Gal* RNAi (BDSC#50680; this stock is often used as a control UAS element to balance the total number of UAS elements), *UAS-Nod-β-gal* (BDSC#9912), UAS-*E-cad* RNAi (BDSC#32904), UAS-*E-cad* RNAi (BDSC#38207). The following stocks were obtained from Vienna Drosophila Resource Center (VDRC): *msps* RNAi (21982), *arl2* RNAi (110627), *sas-4* RNAi (106051). *msps* RNAi knockdown efficiency in larval brains was verified by immunostaining of anti-Msps antibody. Various RNAi knockdown or overexpression constructs were induced using *grh*-Gal4 or *insc*-Gal4 unless otherwise stated.

All experiments were carried out at 25°C, except for RNAi knockdown or overexpression at 29°C, unless otherwise indicated.

### EdU (5-ethynyl-2′-deoxyuridine) incorporation assay

Larvae of various genotypes were fed with food supplemented with 0.2 mM EdU from Click-iT® EdU Imaging Kits (Invitrogen) for 4 h. Larval brains were dissected in PBS and fixed with 4% EM-grade formaldehyde in PBS for 22 min, following by three washes with 0.3% PBST, each wash for 10 min and blocked with 3% BSA in PBST for 30 min. Detection of incorporated EdU by Alexa Fluor azide was according to the Click-iT EdU protocol (Invitrogen). The brains were rinsed twice and subjected to standard immunohistochemistry.

### Immunohistochemistry

*Drosophila* larvae were dissected in PBS, and larval brains were fixed in 4% EM-grade formaldehyde in PBT (PBS + 0.3% Triton-100) for 22 min. The samples were processed for immunostaining as previously described (Li et al., 2017). For α-tubulin immunohistochemistry, larvae were dissected in Shield and Sang M3 medium (Sigma-Aldrich) supplemented with 10% FBS, followed by fixation in 10% formaldehyde in Testis buffer (183 mM KCl, 47 mM NaCl, 10 mM Tris-HCl, and 1 mM EDTA, pH 6.8) supplemented with 0.01% Triton X-100.) The fixed brains were washed once in PBS and twice in 0.1% Triton X-100 in PBS. Images were taken from LSM710 confocal microscope system (Axio Observer Z1; ZEISS), using a Plan-Apochromat 40×/1.3 NA oil differential interference contrast objective, and brightness and contrast were adjusted by Photoshop CS6.

Primary antibodies used in this paper were guinea pig anti-Dpn (1:1000), mouse anti-Mira (1:50, F. Matsuzaki), rabbit anti-Mira (1:500, W. Chia), rabbit anti-GFP (1:3,000; F. Yu), mouse anti-GFP (1:5,000; F. Yu), guinea pig anti-Asl (1:200, C. Gonzalez), rabbit anti-Sas-4 (1:100, J. Raff), mouse anti-α-tubulin (1:200, Sigma, Cat#: T6199), mouse anti-γ-tubulin (1:200, Sigma, Cat#: T5326), rabbit anti-CNN (1:5000, E. Schejter and T. Megraw), rabbit anti-Msps (1:500), rabbit anti-Msps (1:1000, J. Raff), rabbit anti-PH3 (1:200, Sigma, Cat#: 06-570), rat anti-E-cadherin (1:20, DCAD2, DSHB), mouse anti-β-Gal (1:1000, Promega, Cat#: Z3781), rabbit anti-β-galactosidase (1:5000, Invitrogen, A-11132), mouse nc82 (1:20, DSHB) and mouse anti-synaptotagmin1 (1:50, DSHB, 3H2 2D7), rabbit anti-sas-4 (1:200, J. Raff), rabbit anti-Ana2 (Wang et al., 2011) (1:50). The secondary antibodies used were conjugated with Alexa Fluor 488, 555 or 647 (Jackson laboratory).

### Tracking of EB1-GFP comets

Larval brains of various genotypes expressing EB1-GFP under *grh*-Gal4 at various time points were dissected in Shield and Sang M3 insect medium (Sigma-Aldrich) supplemented with 10% FBS. Larval brain explant culture were supplied with fat body from wild-type third instar and live imaging of larval brains were performed with LSM710 confocal microscope system using 40X Oil lens and Zoom factor 6. Larval brains were imaged for 151 seconds with 83 frames acquired for each movie and images were analyzed with NIH ImageJ software. Velocity and kymograph were calculated or generated by KymoButler (Jakobs et al., 2019).

### Quantification and statistical analysis

*Drosophila* larval brains from various genotypes were placed dorsal side up on confocal slides. The confocal z-stacks were taken from the surface to the deep layers of the larval brains (20-30 slides per z-stack with 2 or 3 µm intervals). For each genotype, at least 5 brain lobes were imaged for z-stacks and Image J or Zen software were used for quantifications. Cell diameter of quiescent NSCs was measured based on *grh*>CD8-GFP or Mira cortical localization in Dpn-positive cells.

Statistical analysis was essentially performed using GraphPad Prism 8. Unpaired two-tail t tests were used for comparison of two sample groups and one-way ANOVA or two-way ANOVA followed by Sidak’s multiple comparisons test for comparison of more than two sample groups. All data are shown as mean ± SD. Statistically nonsignificant (ns) denotes P > 0.05, * denotes P <0.05, ** denotes P <0.01, *** denotes P <0.001, and **** denotes P < 0.0001. All experiments were performed with a minimum of two repeats. In general, n refers to number of NSCs counted, unless otherwise indicated.

## Supplemental Figure Legends

**Figure S1.**
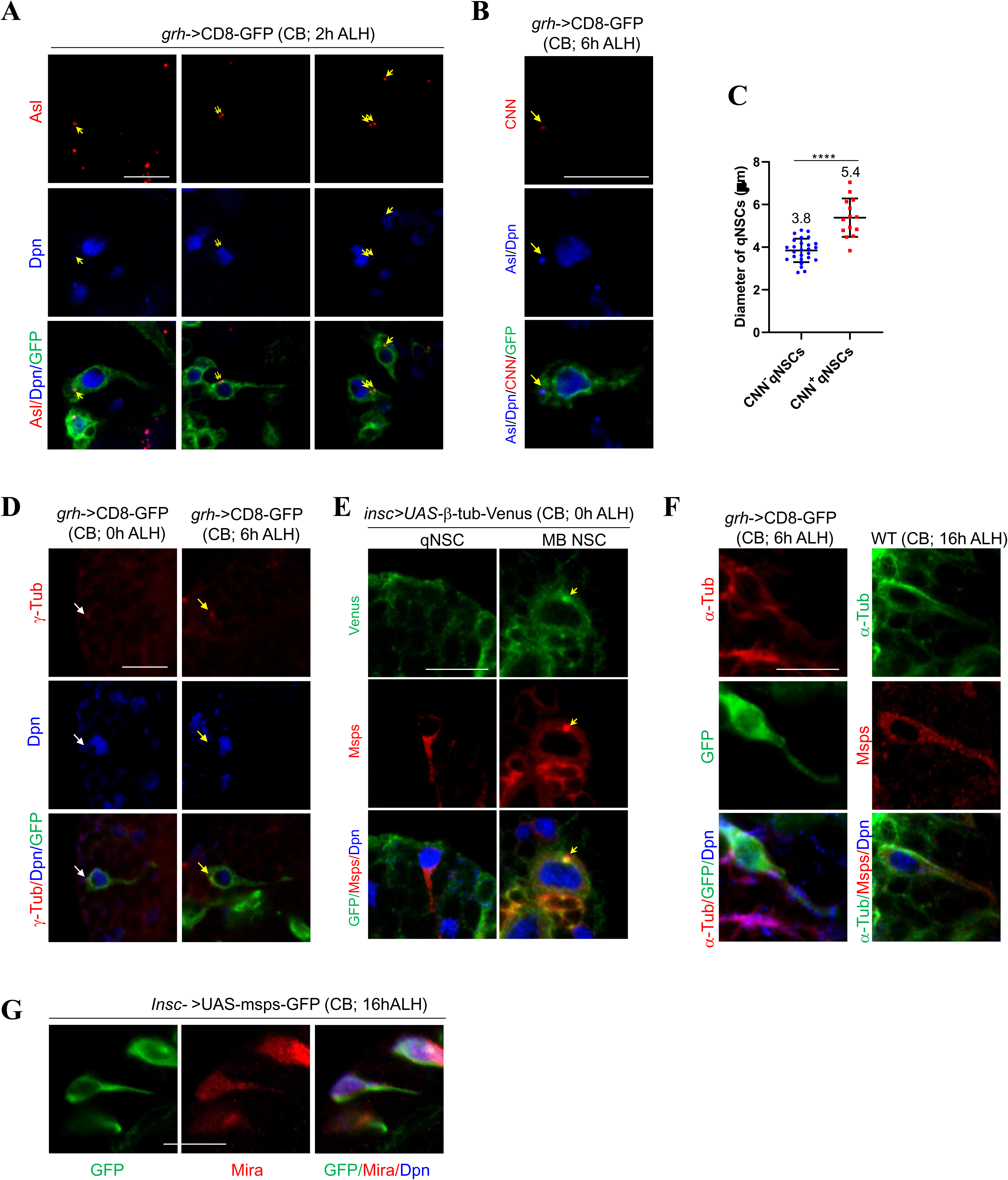
The centrosomes in qNSCs are immature. A) Larval brains at 2h After Laval Hatching (ALH) from *grainy head* (*grh)*-Gal4; UAS-CD8-GFP were labelled with Asterless (Asl), Deadpan (Dpn) and GFP. B) Wild-type larval brains expressing *grh*>CD8-GFP at 6h ALH were labelled with CNN, Asl, Dpn, and GFP. Arrows indicate the centrosomes. C) Quantification of cell diameter of CNN-negative *vs* CNN-positive qNSCs from wild-type brains expressing *grh*>CD8-GFP at 6h ALH. ****p<0.0001. D) Wild-type larval brains expressing *grh*>CD8-GFP at 0h ALH and 6h ALH were labelled with γ-tubulin, Dpn, and GFP. Arrows indicate the centrosome. E) Larval brains at 0h ALH from *insc*-Gal4; UAS-β-tubulin-Venus were labeled with GFP, Msps and Dpn. Quiescent NSCs at the CB were shown. F) Wild-type larval brains expressing *grh*>CD8-GFP at 6h ALH were labelled with α-tubulin, Dpn and GFP and wild-type larval brains at 16h ALH were labelled with α-tubulin, Msps and Dpn. G) Larval brains expressing Msps-GFP under the control of *insc*-Gal4 at 16h ALH were labelled with GFP, Dpn, and Mira. Scale bars: 10 µm.

**Figure S2.**
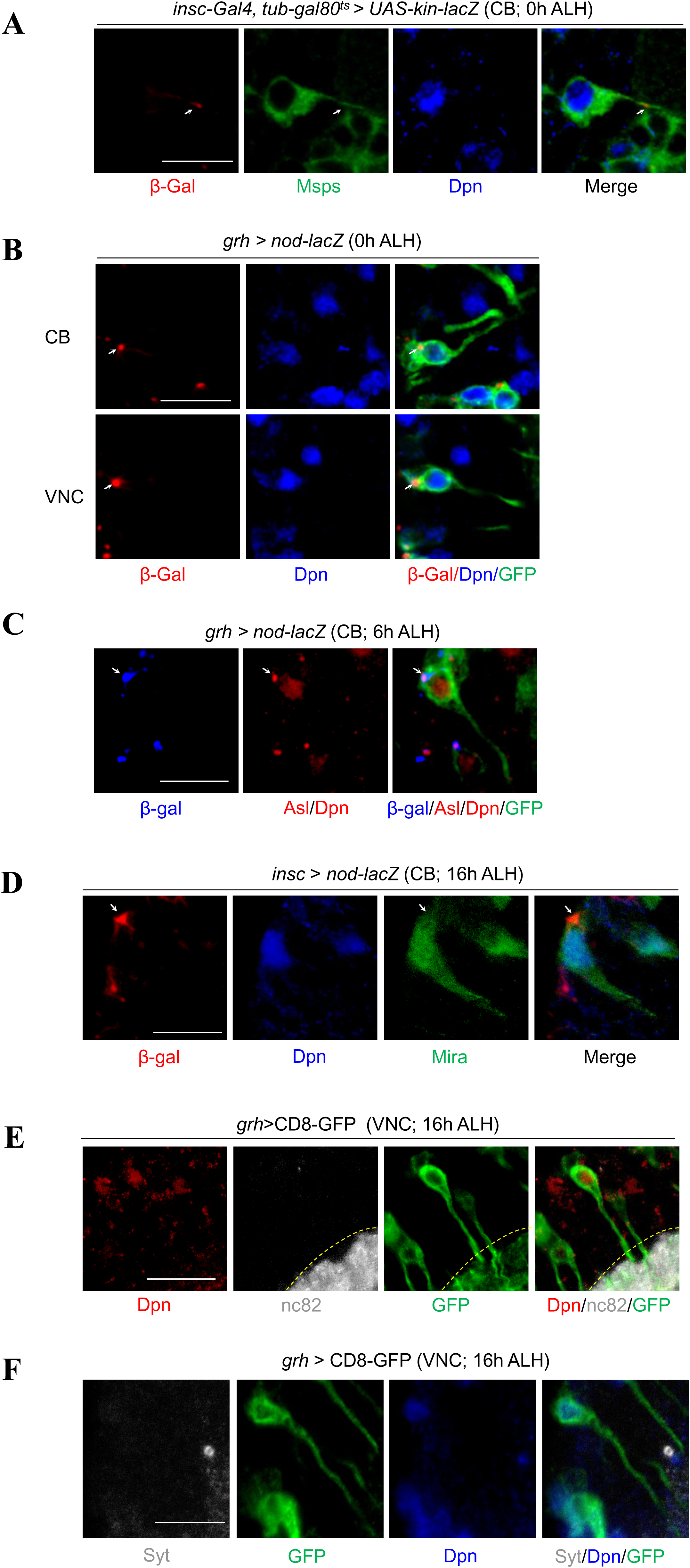
The primary protrusion of qNSCs organizes plus-end-out microtubules. A) Larval brains at 0h ALH, in which *kin-lacZ* was expressed under the control of *insc-*Gal4*, tub-*Gal80^ts^, were labelled with β-Gal, Msps, and Dpn. The primary protrusion of qNSCs were marked by Msps. Quiescent NSCs at the CB were shown. B) Larval brains at 0h ALH from *grh*-Gal4; UAS-nod-lacZ were labelled with β-gal, Dpn, and GFP. Both central brain (CB) and ventral nerve cord (VNC) were shown. C) Larval brains at 6h ALH from *grh*-Gal4; UAS-nod-lacZ were labelled with β-gal, Asl, Dpn, and GFP. Quiescent NSCs at the CB were shown. D) Larval brains at 16h ALH from *insc*-Gal4>*UAS-nod-lacZ* were labelled with β-gal, Dpn, and Mira. Quiescent NSCs at the CB were shown. E) Larval brains at 16h ALH from *grh*-Gal4; UAS-CD8-GFP were labelled with nc82, Dpn, and GFP. Nc82 is an antibody that recognizes Bruchpilot at presynaptic sites. Quiescent NSCs at the VNC were shown. F) Larval brains at 16h ALH from *grh*-Gal4>UAS-CD8-GFP were labelled with Synaptotagmin (Syt; A synaptic marker), Dpn, and GFP. Quiescent NSCs at the VNC were shown. Scale bars: 10 µm.

**Figure S3.**
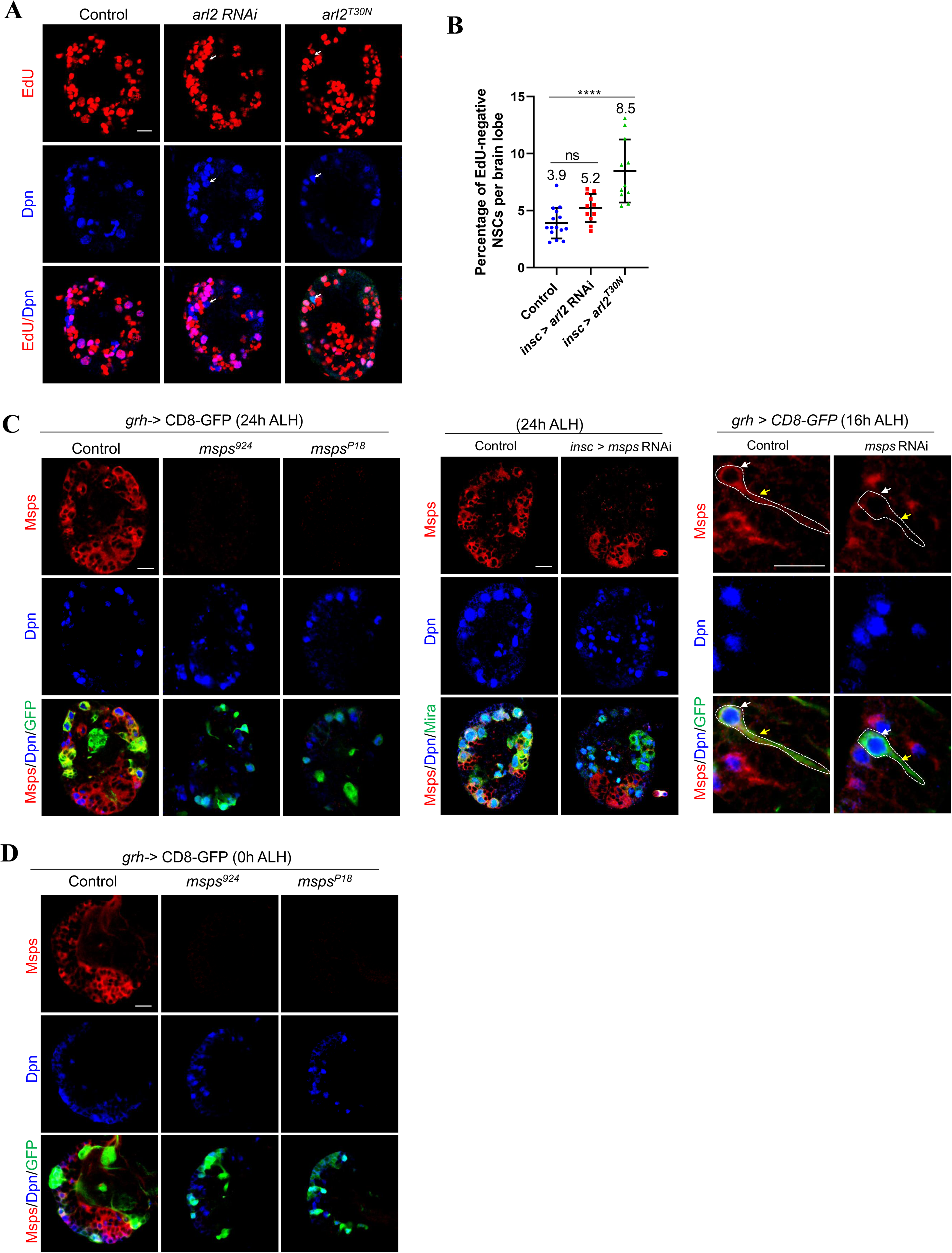
Msps, but not Arl2, is essential for NSC reactivation. A) Larval brains at 24h ALH from control (*insc*-Gal4; *UAS-dicer2 / UAS-β-Gal* RNAi), *arl2* RNAi (VDRC#110627); *UAS-dicer2* and *UAS*-*arl2T30N* under the control of *insc*-Gal4 were analyzed for EdU incorporation. NSCs were marked by Dpn and Mira. B) Quantification of EdU-negative NSCs per brain lobe for genotypes in (a). ****p<0.0001; p=0.1547 (ns). C) Larval brains from various genotypes were labelled with Msps, Dpn and GFP at indicated time points. Left panels, wild-type, *msps^924^*, and *msps^P18^*, all expressing CD8-GFP under the control of *grh*-Gal4. Middle panels, control (*UAS-β-Gal* RNAi) and *msps* RNAi expressing CD8-GFP under the control of *insc*-Gal4. Right panels, control (*UAS-β-Gal* RNAi) and *msps* RNAi under *grh*-Gal4. Note that not all qNSCs were labelled by *grh*>CD8-GFP at early larval stages. D) Larval brains at 0h ALH from wild-type, *msps^924^*, and *msps^P18^* expressing *grh*>CD8-GFP were labelled with Msps, Dpn and GFP. Scale bars: 10 µm.

**Figure S4.**
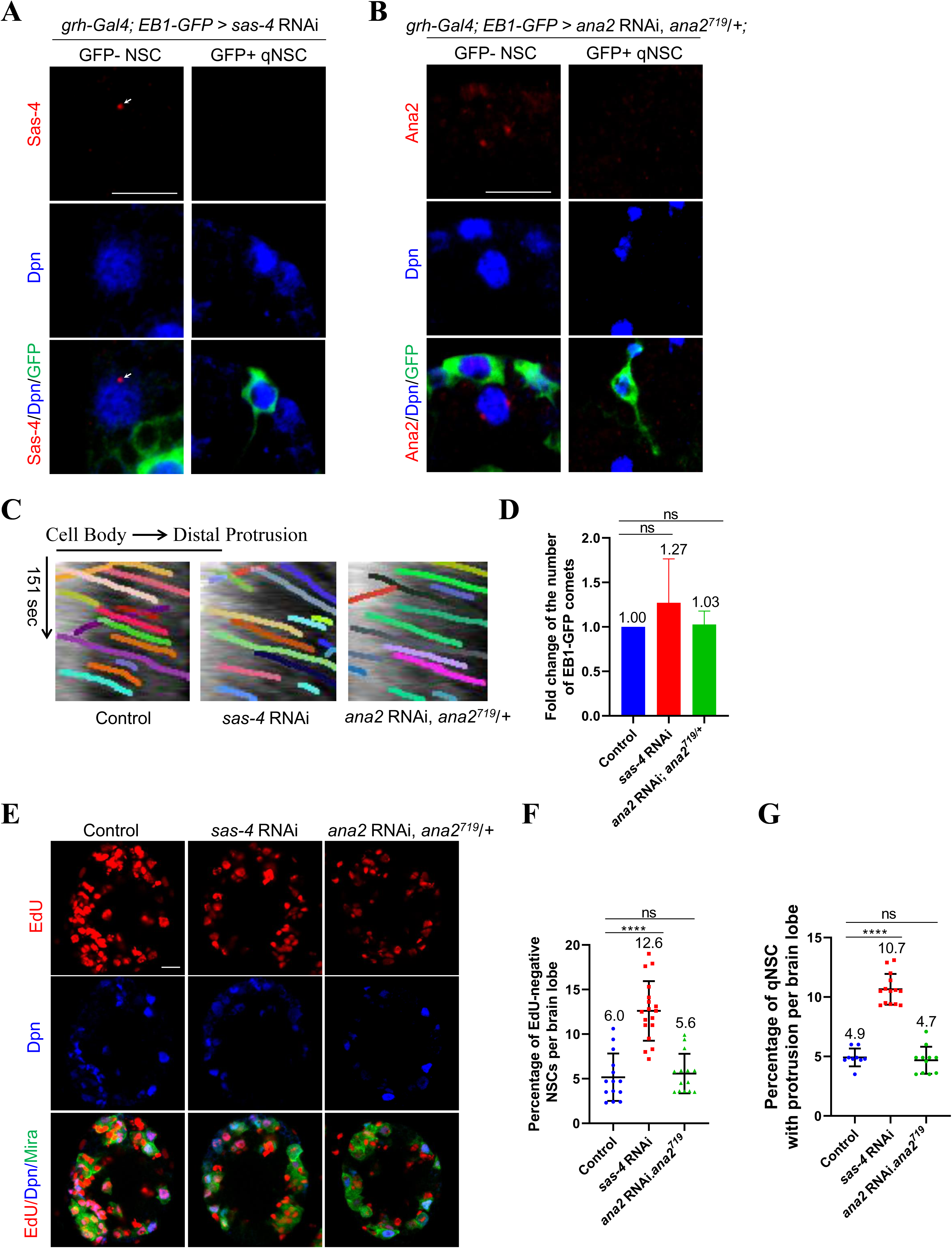
Microtubule assembly in the primary protrusion of qNSCs is centrosome-independent. A) Larval brains at 6h ALH from *sas-4* RNAi with *grh-*Gal4; *UAS-EB1-GFP* were stained with Sas-4, Dpn and GFP. Primary protrusion of qNSCs were marked by GFP. B) Larval brains at 6h ALH *ana2* RNAi *ana2^719^/+* with *grh-*Gal4; *UAS-EB1-GFP* were stained with Ana2, Dpn and GFP. Primary protrusion of qNSCs were marked by GFP. Efficient knockdown of *sas4* or *ana2* in GFP-positive qNSCs with primary protrusion were shown in A-B. Sas4 and Ana2 staining were present in some of the GFP-negative NSCs, that were shown as positive control in A-B. Arrows indicate the centrosome. C) Kymograph of EB-GFP comets movement in the primary protrusion of qNSCs from control (*UAS-β-Gal* RNAi), *sas-4* RNAi (VDRC#106051) and *ana2* RNAi; *ana2^719^/+* with *grh-*Gal4*; UAS-EB1-GFP* at 6h ALH, the horizontal arrow indicates anterograde movement direction from cell body to the tip of the primary protrusion in qNSCs. D) Quantification graph of fold change of the number of EB1-GFP comets in the primary protrusion of qNSCs from genotypes in C). E) Larval brains at 24h ALH from control (*grh*-Gal4; *UAS-dicer2*/*UAS-β-Gal* RNAi) and *sas-4* RNAi (VDRC#106051) and *ana2* RNAi; *ana2^719^/+* controlled under *grh-*Gal4 were analyzed for EdU incorporation. NSCs were marked by Dpn and Mira. F) Quantification graph of EdU-negative NSCs per brain lobe for genotypes in (E). ****p< 0.0001; p=0.9106 (ns). Control, 6.0%, n=14; *sas-4* RNAi, 12.6%, n=18. *ana2* RNAi, *ana2^719^*/+, 5.6%, n=14. G) Quantification graph of percentage of qNSCs with primary protrusion per brain lobe for genotypes in (E). Control, 4.9%, n=9; *sas-4* RNAi, 10.7%, n=14; *ana2* RNAi, *ana2^719^*/+, 4.7%, n=11. ****p< 0.0001; p=0.8681 (ns). Scale bars: 10 µm.

**Figure S5.**
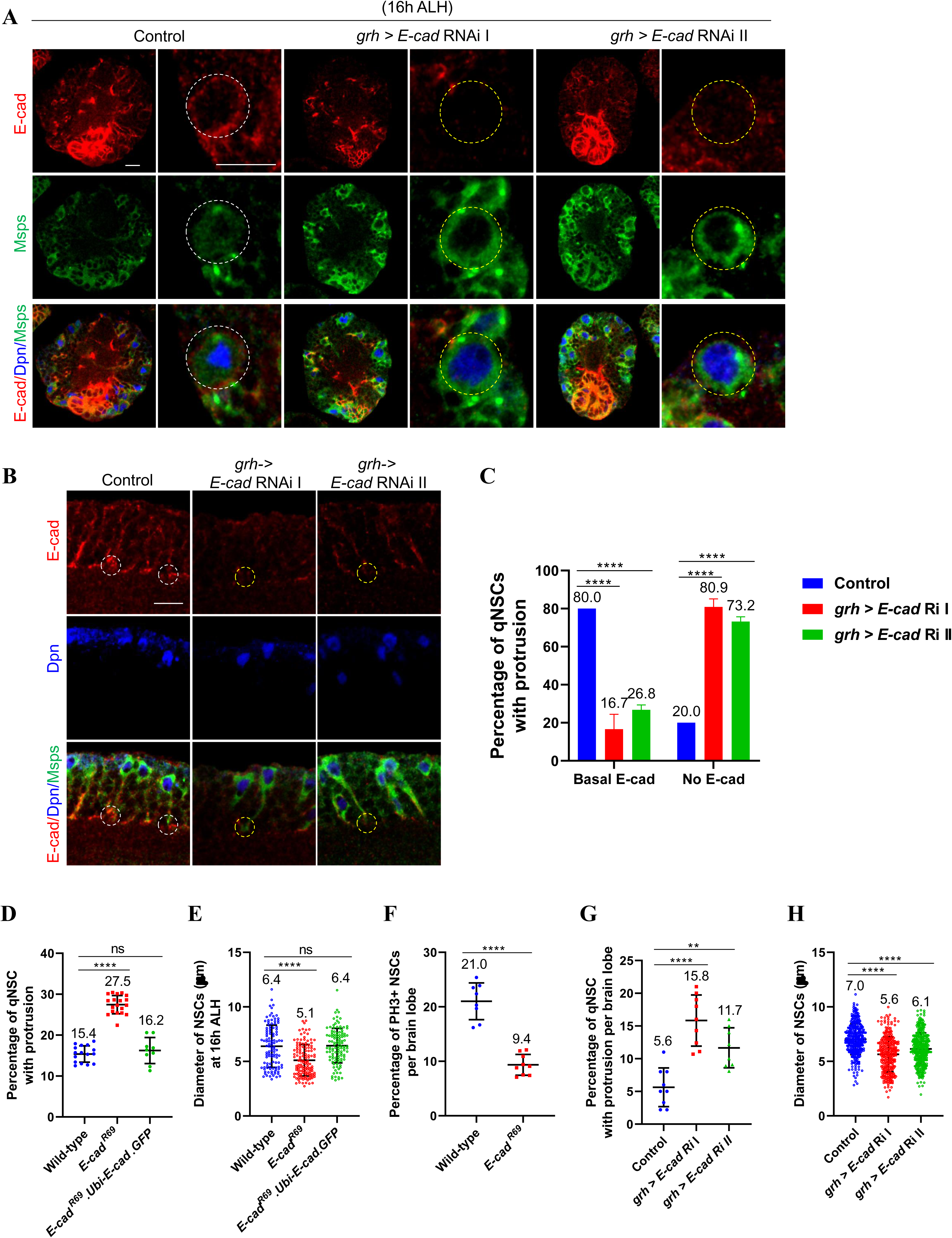
E-cad is dramatically reduced in NSC-neuropil contact upon *E-cad* RNAi knockdown in NSCs. A) Larval brains at 16h ALH from control (*grh*-Gal4; *UAS-Dicer2/UAS-β-Gal* RNAi*), E-cad* RNAi I, and *E-cad* RNAi II under control of *grh*-Gal4; *UAS-Dicer2* were labelled with E-cad, Dpn, and Msps. B) Larval brains (VNC) at 16h ALH from control (*grh*-Gal4; *UAS-Dicer2/UAS-β-Gal* RNAi*), E-cad* RNAi I, and *E-cad* RNAi II under control of *grh*-Gal4; *UAS-Dicer2* were labelled with E-cad, Dpn, and Msps. Primary protrusion of qNSCs were marked by Msps.C) Quantification of E-Cadherin basal localization at putative NSC-neuropil contact sites in qNSCs from genotypes in (b). “No E-cad” means absent or strongly reduced E-cad observed at NSC-neuropil contact sites. ****p<0.0001. D) Quantification graph of percentage of qNSCs with primary protrusion in wild-type, *E-cad^R69^* and *E-cad^R69^.Ubi-p63E-E-cad.GFP* at 16h ALH. ****p<0.0001; p=0.6339 (ns). E) The protrusion diameter of the cell body in NSCs at 16h ALH from wild-type, *E-cad^R69^* and *E-cad^R69^.Ubi-p63E-E-cad.GFP.* NSCs were marked by Dpn and Mira. ****p<0.0001; p=0.9658 (ns). F) Quantification graph of percentage of NSCs with PH3 per brain lobe from Wild-type and *E-cad^R69^* at 16h ALH. ****p< 0.0001. G)) Quantification graph of percentage of qNSCs with primary protrusion in control, *E-cad* RNAi I, and *E-cad* RNAi II under *grh*-Gal4; *UAS-Dicer2.* The protrusion was labelled by Mira. ****p<0.0001; **p=0.0022. H) Quantification graph of diameter of the cell body in NSCs at 24h ALH from control, E-cad RNAi I, and E-cad RNAi II under control of grh-Gal4; UAS-Dicer2. NSCs were marked by Dpn and Mira. ****p<0.0001. Scale bars: 10 µm.

**Movie S1.** Time-lapse imaging of EB1-GFP comets in the primary protrusion of qNSCs in larval brains at 6h ALH from *grh*-Gal4; *UAS-EB1-GFP*. Time scale: minute: second. Scale bar: 10 µm.

**Movie S2**. Time-lapse imaging of EB1-GFP comets in the primary protrusion of qNSCs in larval brains at 2h ALH from *grh*-Gal4; *UAS-EB1-GFP*. Time scale: minute: second. Scale bar: 10 µm.

**Movie S3.** Time-lapse imaging of EB1-GFP comets in the primary protrusion of qNSCs in larval brains at 6h ALH from *grh*-Gal4; *msps^P18^ UAS-EB1-GFP*. Time scale: minute: second. Scale bar: 10 µm.

**Movie S4.** Time-lapse imaging of EB1-GFP comets in the primary protrusion of qNSCs in larval brains at 6h ALH from *grh*-Gal4; *msps^P18/P^ UAS-EB1-GFP*. Scale bar: 10 µm.

**Movie S5.** Time-lapse imaging of EB1-GFP comets in the primary protrusion of qNSCs in larval brains at 6h ALH from sas-4 RNAi with *grh*-Gal4; *UAS-EB1-GFP*. Scale bar: 10 µm.

**Movie S6.** Time-lapse imaging of EB1-GFP comets in the primary protrusion of qNSCs in larval brains at 6h ALH from *ana2* RNAi, *ana2^719^* with *grh*-Gal4; *UAS-EB1-GFP*. Scale bar: 10 µm.

